# PopAmaranth: A population genetic genome browser for grain amaranths and their wild relatives

**DOI:** 10.1101/2020.12.09.415331

**Authors:** José Gonçalves-Dias, Markus G Stetter

**Author notes:** Dept. for Plant Sciences, University of Cologne, Cologne, Germany (Markus G Stetter).

## Abstract

The last decades of genomic, physiological, and population genetic research have accelerated the understanding and improvement of a numerous crops. The transfer of methods to minor crops could accelerate their improvement if knowledge is effectively shared between disciplines. Grain amaranth is an ancient nutritious pseudocereal from the Americas that is regaining importance due to its high protein content and favorable amino acid and micronutrient composition. To effectively combine genomic and population genetic information with molecular genetics, plant physiology, and use it for interdisciplinary research and crop improvement, an intuitive interaction for scientists across disciplines is essential. Here, we present PopAmaranth, a population genetic genome browser, which provides an accessible representation of the genetic variation of the three grain amaranth species (*A. hypochondriacus, A. cruentus*, and *A. caudatus*) and two wild relatives (*A. hybridus* and *A. quitensis*) along the *A. hypochondriacus* reference sequence. We performed population-scale diversity and selection analysis from whole-genome sequencing data of 88 curated genetically and taxonomically unambiguously classified accessions. We incorporate the domestication history of the three grain amaranths to make an evolutionary perspective for candidate genes and regions available. We employ the platform to show that genetic diversity in the water stress-related MIF1 gene declined during amaranth domestication and provide evidence for convergent saponin reduction between amaranth and quinoa. These examples show that our tool enables the detailed study of individual genes, provides target regions for breeding efforts and can enhance the interdisciplinary integration of population genomic findings across species. PopAmaranth is available through amaranthGDB at amaranthgdb.org/popamaranth.html

**Significance:** Sharing population genetic results between disciplines can facilitate interdisciplinary research and accelerate the improvement of crops. Since the onset of genome sequencing online genome browser platforms have provide access to features of an organisms genetic information. Rarely this has been extended to population-wide summary statistics for evolutionary hypothesis testing. We implemented a population genetic genome browser PopAmaranth for three grain amaranth species and their two wild relatives. The intuitive and user-friendly interface of PopAmaranth makes the genetic diversity of the species complex available to broad audience of biologists across disciplines. We show how our tool can be used to study convergence across distant genera and find signals of past selection in domestication and stress related genes. Community platforms and genome browsers are an integrative element of numerous study systems. PopAmaranth can serve as template for other research communities to integrate and share their results.

## Main

Genome sequencing, genome-assisted breeding, and molecular breeding techniques have accelerated the improvement of numerous major crops (Wallace *et al.* 2018; Lemmon *et al.* 2018). The availability of genome-wide diversity data of crops and their wild relatives has allowed to identify and study candidate genes of agronomic significance (Hufford *et al.* 2012; Huang *et al*. 2012; Wang *et al.* 2020). These candidates can then be validated through molecular genetics (Ross-Ibarra *et al.* 2007; Fernie and Yan 2019; Sedeek *et al.* 2019; Wang *et al.* 2020). To facilitate the interdisciplinary use of population genetic results, it is essential to provide summary statistics in an intuitive and user-friendly way.

Different platforms have been developed to make genomic resources available across disciplines and have enabled the integration of complementary research areas (Lawrence *et al.* 2004; Alonso-Blanco *et al.* 2016; Jin *et al.* 2013). Online genome browser platforms such as Ensemble (Bolser *et al.* 2016) and Phytozome (Goodstein *et al.* 2012) have become a standard interface to interact with genome sequences and annotations and are used across research fields. Genome browsers provide access to reference genome sequences and gene annotations for numerous plant species but most browsers only provide data for a single reference individual per species. Species-specific browsers include sequence data and variant calls for a large number of individuals (e.g., Lawrence *et al.* 2004; Dash *et al.* 2016; Krishnakumar *et al*. 2015; Mansueto *et al.* 2017; Kudo *et al.* 2017), but do not allow a direct inference of a population scale genome-wide diversity across related species. For few non-plant model species population genetic genome browsers, providing population-scale summary statistics have been developed (Casillas *et al.* 2018). For plant and crop species, in particular minor crops, such resources are currently unavailable.

Novel and under-utilized crops have a high potential to contribute to sustainable food production, as many such crops are tolerant to abiotic and biotic factors and are of high nutritional value (Mayes *et al.* 2012). Amaranth is an under-utilized crop that has been cultivated for its grains as pseudo-cereal and its edible leaves as a vegetable (Sauer 1967; Joshi *et al.* 2018). Three grain amaranth species, *Amaranthus caudatus, A. cruentus* L., and *A. hypochondriacus* L., have been domesticated for their grain from a common wild ancestor, *A. hybridus* L. (Stetter *et al.* 2020). Another wild relative, *A. quitensis* Kunth, is suspected to be involved in the domestication of the South American *A. caudatus*, although its role and contribution to the crop remain unclear (Stetter *et al.* 2017, 2020). The repeated domestication of amaranth presents an interesting model to study genetic parallelisms along selection gradients, and the combination of genomics, quantitative genetics, and molecular dissection of gene function has a high potential to improve grain amaranth.

First resources that allow the functional study of traits have been developed for amaranth. On the one hand, numerous genomic resources, including a high-quality reference genome (Lightfoot *et al.* 2017) and a transcriptome (Clouse *et al.* 2016), genome-wide marker data (Mallory *et al.* 2008; Stetter *et al.* 2017, 2020) and QTL regions for different traits (Lightfoot *et al.* 2017; Stetter *et al.* 2020) have been identified. On the other hand, a number of molecular methods have been adapted for the crop, including molecular gene function identification (Massange-Sanchez *et al.* 2016), state-of-the-art transient ‘hairy’ roots expression systems (Castellanos-Arévalo *et al.* 2020), and stress physiology assays (Parra-Cota *et al.* 2014; Massange-Sanchez *et al.* 2015). Combined, these resources can elevate amaranth research and improvement if results and data are available and accessible for researchers across disciplines.

Here, we present PopAmaranth, an interactive genome-wide population genetic browser for amaranth. PopAmaranth facilitates browsing a number of population genetic summary statistics and selection signals, gene annotation, and variant calls of the three grain amaranths and two wild relatives along the amaranth genome. We defined a curated set of 88 morphologically and genetically identified samples with whole-genome sequencing data to represent the five populations. Currently, PopAmaranth provides three categories of summary statistics, namely genetic diversity, population differentiation, and selection signals, plus variant calls and annotation tracks, in a total of more than 40 tracks. We show how the tool allows a user-friendly way to screen evolutionary signals for candidate genes and compare them between populations by identifying selection signals in a stress gene previously identified in one of the grain amaranths and in an ortholog quinoa domestication gene that shows convergent signals of selection in amaranth. PopA-maranth is embedded in amaranthGDB and is accessible from amaranthgdb.org/popamaranth.html.

## Methods

### Data and filtering

We used whole-genome sequencing data of 116 accession from five amaranth species, including the three grain amaranths (24 *A. hypochondriacus*, 24 *A. cruentus*, and 34 *A. caudatus* samples) and their two wild relatives, 9 *A. hybridus* and 25 *A. quitensis* (Stetter *et al.* 2020, Table S1). The sequencing reads were aligned to the *A. hypochondriacus* reference sequence V 2.0 (Lightfoot *et al*. 2017).

We performed principal component analysis (PCA) on the full set of accessions to remove individuals with ambiguous species clustering using PCAngsd (Meisner and Albrechtsen 2018) and prcomp and autoplot functions in R. We excluded samples that did not cluster with the morphologically designated species according to their passport data (Figure S3).

We calculated the site allele frequency likelihood based on individual genotype likelihoods for each of the five species using the -doSaf 1 function on ANGSD (Korneliussen *et al.* 2014). We removed sites with a minimum map quality below 30, minimum base qscore below 20, and a flagstat (Li *et al.* 2009) above 255, keeping only primary reads (-doSaf 1, -GL 2, -remove_bads 1, -minMapQ 30. -minQ 20). In addition, we removed all sites with more than 66% missing values (-minInd=1/3*n).

### Population genetic browser tracks

Using realSFS saf2theta functions on ANGSD, we calculated the folded site frequency spectrum and estimated per site thetas (population scaled mutation rate). Consequently, we calculated nucleotide diversity (*π*) and Wu and Watterson estimator (*θ*) in non-overlapping windows of 5000 bp. We only kept windows with more than 30% of the sites called in a given window. We used the scikit-allel python library (https://doi.org/10.5281/zenodo.597309l) to calculate per site heterozygosity statistics (*H*_*exp*_,*H*_*obs*_, and *F*) for each of the five populations after sub-setting variant calls (VCF file) from Stetter *et al*. (2020) to include the samples described above using VCFtools (Danecek *et al.* 2011). A yellow horizontal line denotes the genome-wide mean for each of the summary statistics. To visually distinguish the deviation from the genome-wide mean, values below the mean are represented in red and above the mean in blue. Further, we indicate the strength of deviation by adding dark gray and light grey shadings for one and two standard deviations from the mean, respectively.

We calculated pairwise Weir-Cockerham F_*st*_ (Wright 1950) as a measure for genetic differentiation for each pair of populations using ANGSD (Korneliussen *et al.* 2014). We used these values as input to calculate pairwise F_*st*_ in non-overlapping windows of 5000bp along the genome.

We applied Raised Accuracy in Sweep Detection (RAiSD) (Alachiotis and Pavlidis 2018) with default setting (20 SNP windows) on the subset VCF data from Stetter *et al*. (2020) to detect signals of selective sweeps within each population. We considered windows on the top 1 % *µ* values as outliers and under positive selection (*A. caudatus*: 17650 windows; *A. cruentus* 16546; *A. hypochondriacus*: 17932 *A. hybridus*: 43415; and *A. quitensis*: 15854). We merged all overlapping windows to create stretches of selective sweeps.

We employed ANGSD (Korneliussen *et al.* 2014) with the parameters described above for *π* and *θ* to calculated Tajima’s D in non-overlapping 5 kb windows.

Using the nucleotide diversity estimated for each of the species, we calculated relative nucleotide diversity. We divided *π* for each of the domesticated species (*A. caudatus, A. cruentus*, and *A. hypochondriacus* by *π* of their wild ancestor, *A. hybridus*. We only used windows where both species had data after filtering for the number of genotyped sites.

### Browser implementation and annotation

We provided access to the summary statistics described above as an interactive tool through JBrowse 1.16.9 (Skinner *et al.* 2009). We added the reference sequence and gene annotation, including exons, intros, CDS, mRNA, and UTRs from Lightfoot *et al*. (2017) available through Phytozome (Goodstein *et al.* 2012). For each summary statistic a color gradient summary plot combining all species was added. Further, we added the “Variant” category, providing variant data for biallelic SNPs within each species from Stetter *et al*. (2020) (not including variants fixed between populations).

### PopAmaranth application to candidate genes

We downloaded the sequence of the water stress-related MIF1 gene reported in Huerta-Ocampo *et al*. (2011) from the NCBI database and used BLASTn (Altschul *et al.* 1990) to identify the gene ID in the *A. hypochondriacus* V2 reference sequence on Phytozome. Using the same procedure, we studied the triterpene saponin biosynthesis activating regulator-1 (TSAR-1) gene from *Chenopodium quinoa* (Jarvis *et al.* 2017).

## Results

### Sample filtering

*Amaranthus* species are difficult to taxonomically classify because of their high morphological similarity (Sauer 1967). Therefore, we sub-sampled the original dataset from Stetter *et al*. (2020) based on the genetic clustering in the PCA and species delimitation in Germplasm Resources Information Network (GRIN). We selected each species according to their clustering in the first three principal components (Figure S3). After filtering, our sample consisted of 88 genetically and morphologically defined samples representing the five species, with 28 individuals classified as *A. caudatus* L., 21 *A. cruentus* L., 18 *A. hypochondriacus* L., 12 *Amaranth quitensis* Kunth, and 9 *A. hybridus* L. (Figure 1 and table S1).

**Figure 1.**
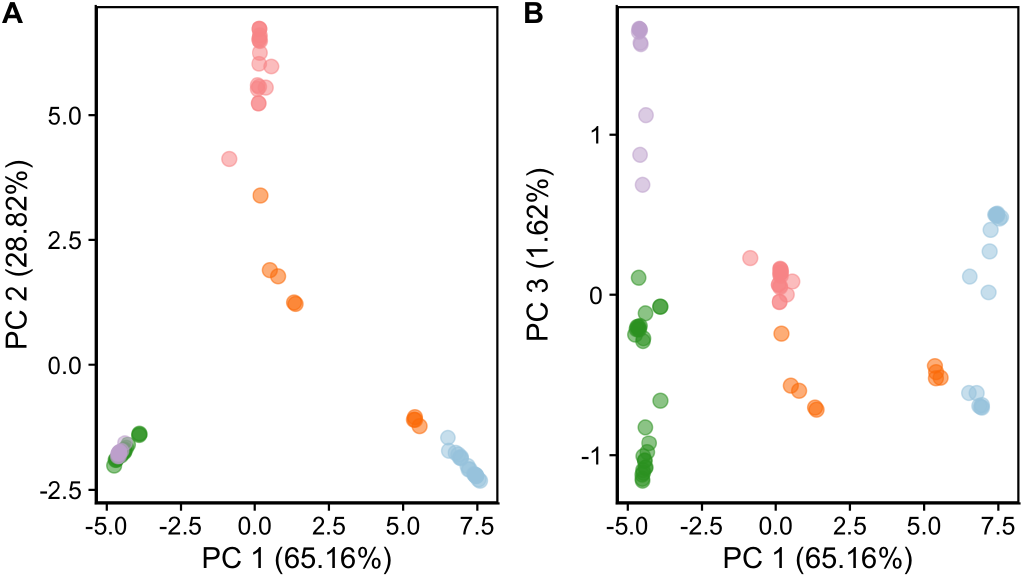
Principal Component Analysis with filtered samples. Each dot represents each of the 88 samples. *A. caudatus* (green), *A. cruentus* (blue), *A. hybridus* (orange), *A. hypochondriacus* (rose), *A. quitensis* (purple). Axis show the percentage of variance explained by each principal component

### Categories and Tracks

We created PopAmaranth relative to the high-quality *A. hypochondriacus* reference genome (Lightfoot *et al*. 2017) and added the gene annotation as functional guide. We calculated nine summary statistics from whole-genome sequencing data for each of the five species. The tracks are grouped into five categories, namely annotation, differentiation, diversity, selection, and variant calls (Table 1 and 1). Each category includes tracks one color gradient summary track combining data of a summary statistic for all species.

**Table 1.** Tracks.

| Track | Description |
| --- | --- |
| **Annotation** |  |
| Reference Genome v2.0 | *Amaranthus hypochondriacus* reference genome v2.0 (Lightfoot *et al.* 2017) |
| Gene Annotation v2.1 | *Amaranthus hypochondriacus* gene annotation with subfeatures, including CDS, mRNA and UTRs |
| **Differentiation** |  |
| $F_{st}$ | Fixation Index, average pairwise differences Weir and Cockerham (1984) |
| **Diversity** |  |
| Wu & Watterson $\theta$ | Estimator of genetic diversity in a population (Watterson 1975) |
| Expected heterozygosity | Expected heterozygosity for a SNP under Hardy-Weinberg equilibrium |
| Observed heterozygosity | Observed heterozygosity for a SNP genotype. |
| Inbreeding coefficient | Inbreeding coefficient (F) for each variant |
| Nucleotide diversity ($\pi$) | Nei's nucleotide diversity (Nei and Li 1979) |
| **Selection** |  |
| Tajima's D | Difference between the mean number of pairwise differences and the number of segregating sites Tajima (1989) |
| Relative nucleotide diversity | Ratio of nucleotide diversity between a domesticated species and their wild ancestor (*A. hybridus*) |
| Selective Sweep (RAiSD ($\mu$)) | $\mu$ statistic for selective sweep detection |
| **Variant Call** |  |
| VCF | Called SNPs with a given species and their genotype frequency |

### Differentiation

Tracks in the differentiation category represent all pairwise F_*st*_ comparisons in 5 kb windows. The genome-wide pairwise F_*st*_ ranged from 0.17 between *A. caudatus* and *A. quitensiss* to 0.68 between *A. caudatus* and *A. cruentus*. As observed before, F_*st*_ between crop species was higher than between the crops and their wild ancestor for *A. caudatus* and *A. hypochondriacus* (Stetter *et al.* 2020). Although, we found higher F_*st*_ between *A. cruentus* and *A. hybridus* (0.69) than between *A. cruentus* and *A. hypochondriacus* (0.57).

### Diversity

Genetic diversity patterns along the genome can give insights into the evolutionary history of a population. Hence, we calculated several diversity statistics along the genome. Inbreeding coefficients and expected and observed heterozygosity are reported on a per-site basis for each SNP that segregated within a population. In addition to SNP-based statistics, we provide windowed diversity measures, including Wu & Watterson *θ* and nucleotide diversity *π* in 5 kb non-overlapping windows. Consistent with previous findings, the three grain amaranths had a lower mean *π* (0.005-0.010) compared to their wild ancestor *A. hybridus* (0.019) (Stetter *et al.* 2020). Wu & Watterson *θ* was also lower for domesticated amaranth species (0.004-0.007) compared to *A. hybridus* (0.023).

### Selection

We calculated three different summary statistics to detect signals of selection along the genome. Tracks displaying Tajima’s D were calculated in 5 kb windows for each species. Tajima’s D was higher for domesticated species (1.443 in *A. caudatus*, 1.773 in *A. cruentus*, and -0.105 in *A. hypochondriacus*) than for their wild ancestor *A. hybridus* (−0.597), indicating a domestication bottleneck. *A. quitensis* had a mean Tajima’s D of 2.037 also suggesting a recent population contraction.

We employed RAiSD to detect signals of selective sweeps in 20 SNP windows within each species. The top 1% of all windows were considered outliers and suggest regions of positive selection. After merging adjacent outliers, we found 973 non-overlapping windows with positive selection signals in *A. caudatus*, 1,096 in *A. cruentus*, 1,121 *A. hypochondriacus*, 2,452 *A. hybridus*, and 1,275 windows in *A. quitensis*. To investigate the signal of domestication-related selection, we added the relative nucleotide diversity between each crop and their wild ancestor *A. hybridus* in 5 kb windows. While the genome-wide *π* was lower for all three crops (see “Diversity”), relative *π* allows to visualize deviations from this genome-wide mean and detect outlier signals in individual regions.

### Variant Calls

Individual variants give access to an individuals’ genotype. Molecular biologists might be interested in evaluating natural alleles of a gene of interest, and plant breeders could use individuals with specific variants to enrich their gene pools. We provide variant data for all five species representing their genotype frequency within the population. Each variant track only displays variants within the given population (not including fixed variants between populations). A total of 4,961,210 variants for *A. caudatus*, 4,075,368 for *A. cruentus*, 4,551,278 for *A. hypochondriacus*, 12,238,589 for *A. hybridus*, and 2,342,505 for *A. quitensis* along the genome are available.

### PopAmaranth case study

To show the utility of PopAmaranth, we evaluated the evolutionary signals for a gene that was molecularly shown to be involved in the response of *A. hypochondriacus* to water stress (Huerta-Ocampo *et al.* 2011). We found that MIF1 (AH-017582) showed lower nucleotide diversity, decreased expected heterozygozity, and a relative nucleotide diversity below the genome-wide average in all three grain amaranth species. Also, we identified a selective sweep in *A. hypochondriacus* around this gene (Figure 2).

**Figure 2.**
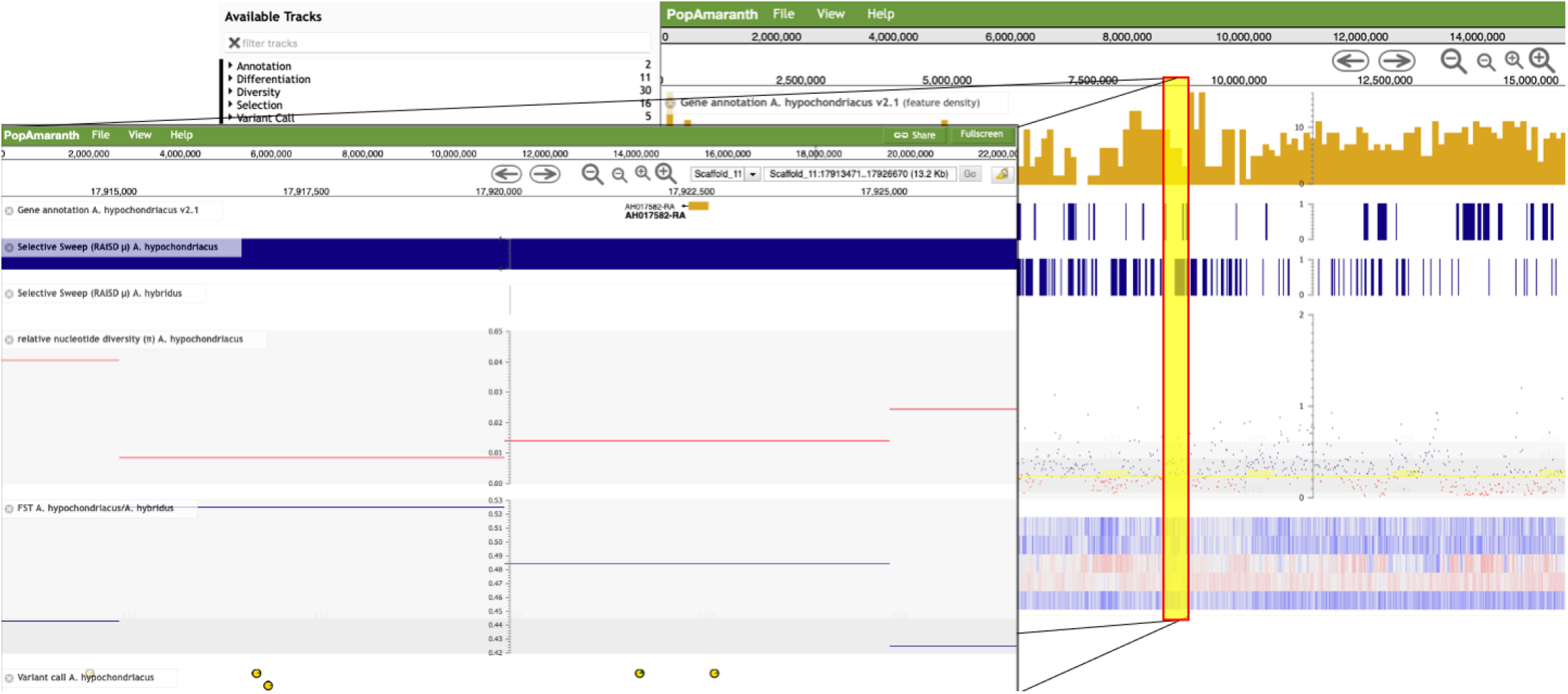
PopAmaranth screen view. Background panel: Zoomed out user view along a chromosome. Search field provides access to genome positions or gene names. Front panel: example is illustrated with a zoom-in region for the water-stress related MIF1 gene (AH-017582).

In addition to the amaranth specific use, PopAmaranth facilitates the evaluation of hypothesis beyond the species. To show its utility to study convergent selection signals across distant families, we evaluated population genetic signals around the amaranth ortholog to the triterpene saponin biosynthesis activating regulator 1 - TSAR1 (AH-019562), a key regulator for seed saponin content in *Chenopodium quinoa* (Jarvis *et al.* 2017). We found signals of selective sweeps in the three grain amaranth species. Furthermore, the relative diversity compared to the wild ancestor was below the genome-wide mean, suggesting selection during amaranth domestication (Figure S4).

## Discussion

Over the last decades, large-scale population genomic data revealed insights into the evolution and adaptation of crops. Providing access to results in a user-friendly and interactive way opens paths to better integrate data from different research areas. Our population genomic genome browser, PopAmaranth, aims to provide such an intuitive tool for amaranth population genetic results. The inclusion of five different species involved in the crop domestication history of facilitates hypothesis testing along this evolutionary gradient.

For other plant species, i.e., maize (Lawrence *et al*. 2004), tomato (Fernandez-Pozo *et al.* 2015), and arabidopsis (Alonso-Blanco *et al*. 2016) accessible platforms of genomic and evolutionary data are integral parts of the research communities. We hope that PopAmaranth and the higher-level framework amaranthGDB will help establish an amaranth community that benefits from the interdisciplinary exchange.

Our results show how PopAmaranth can be employed to add an evolutionary perspective to different molecular questions. We identify previously unknown signals of selection in stress-related MIF1 gene, which might have been under selection during amaranth domestication. In most crops, domestication led to a reduction in stress resilience compared to their wild ancestors. Hence, the reduction in diversity might represent selection against the tolerant allele to free resources for increased crop productivity (Koziol *et al.* 2012). Our browser allows the selection of genotypes with different alleles within grain amaranths and in wild amaranth, enabling the identification of stress-tolerance alleles and potentially the reintroduction of such alleles into breeding programs.

On a broader scale, PopAmaranth also facilitates the comparison of convergent adaptation signals between more distant taxa. For instance, our finding of convergent selection between quinoa and amaranth in a saponin-related gene suggests that in both quinoa and amaranth the saponin content was reduced to improve the palatability of the grains (Jarvis *et al.* 2017). Saponins confer toxicity to protect wild plants against birds but reduce the nutritional quality of seeds for human consumption and animal feed (Oleszek *et al.* 1999; Mroczek 2015). Hence, our platform allowed to identify the convergent selection between the two pseudocereals, demonstrating its utility to evaluate selection signals across taxa. This is of particular use for close relatives of weedy *Amaranthus* species that are of evolutionary and agronomic interest and have been aligned to the same reference genome used in our browser (Montgomery *et al.* 2020).

We aimed for a generalized usage of diversity and differentiation estimates. Therefore, we only selected unambiguous samples of each species, base on morphological and genetic classifications. A clear grouping is crucial for a reference tool, as misclassified samples would confound population-wide signals (Rieseberg and Wendel 2004). Our sub-sampling approach is conservative regarding genetic diversity, as it excludes more differentiated individuals from the analysis. Reported values of genetic differentiation (F_*st*_) between species could be inflated due to the lack of intermediate individuals. The increased differentiation by sub-sampling potentially led to the higher F_*st*_ value between *A. cruentus* and *A. hybridus* compared to previous results (Stetter *et al.* 2020). While there is a trade-off between including additional individuals and the potential for undiscovered diversity, our goal was a defined and distinguished set of samples representing each species. The inclusion of only core individuals of each species further allows the comparison and classification of less distinct individuals using our set.

Altogether, we incorporated a well-defined set of individuals with congruent data filtering to estimate population-wide diversity statistics for the three grain amaranth species and two wild relatives. The identification of selection signals in candidate genes within amaranth and beyond shows the utility of the browser for a range of researchers. PopAmaranth and the amaranthGDB platform will help build and grow the amaranth research community and facilitate interdisciplinary research to ultimately improve the crop.

## Availability

PopAmaranth is available at https://amaranthgdb.org/popamaranth.html. A static version can be found at: 10.6084/m9.figshare.13340798 Code is available at https://github.com/cropevolution/PopAmaranth

## Acknowledgments

We thank the RRZ team at University of Cologne, for hosting PopAmaranth, the de Meaux lab for testing and feedback on the browser, and all members of the Stetter Lab for discussion and suggestions. We acknowledge the support of the Deutsche Forschungsgemeinschaft under Germany’s Excellence Strategy – EXC-2048/1 – Project ID 390686111 to MGS.

## Supplement

**Figure S3.**
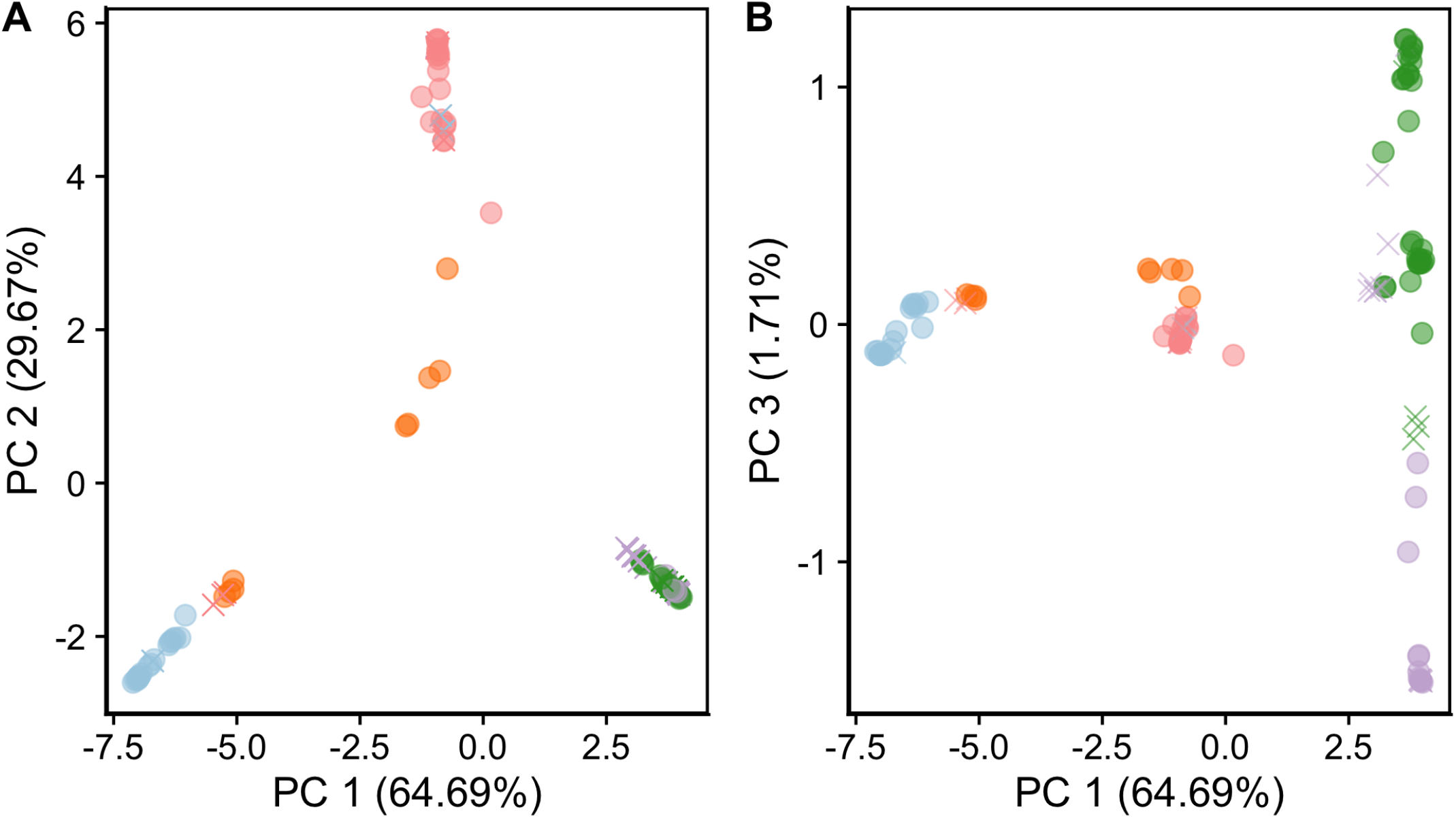
PCA before filtering. Each symbol represents each of the 116 sample. Circles represent amaranth samples included in the study. Removed samples are marked with crosses. *A. caudatus* (green), *A. cruentus* (blue), *A. hybridus* (orange), *A. hypochondriacus* (rose), *A. quitensis* (purple). Axis show the percentage of variance explained by each principal component.

**Figure S4.**
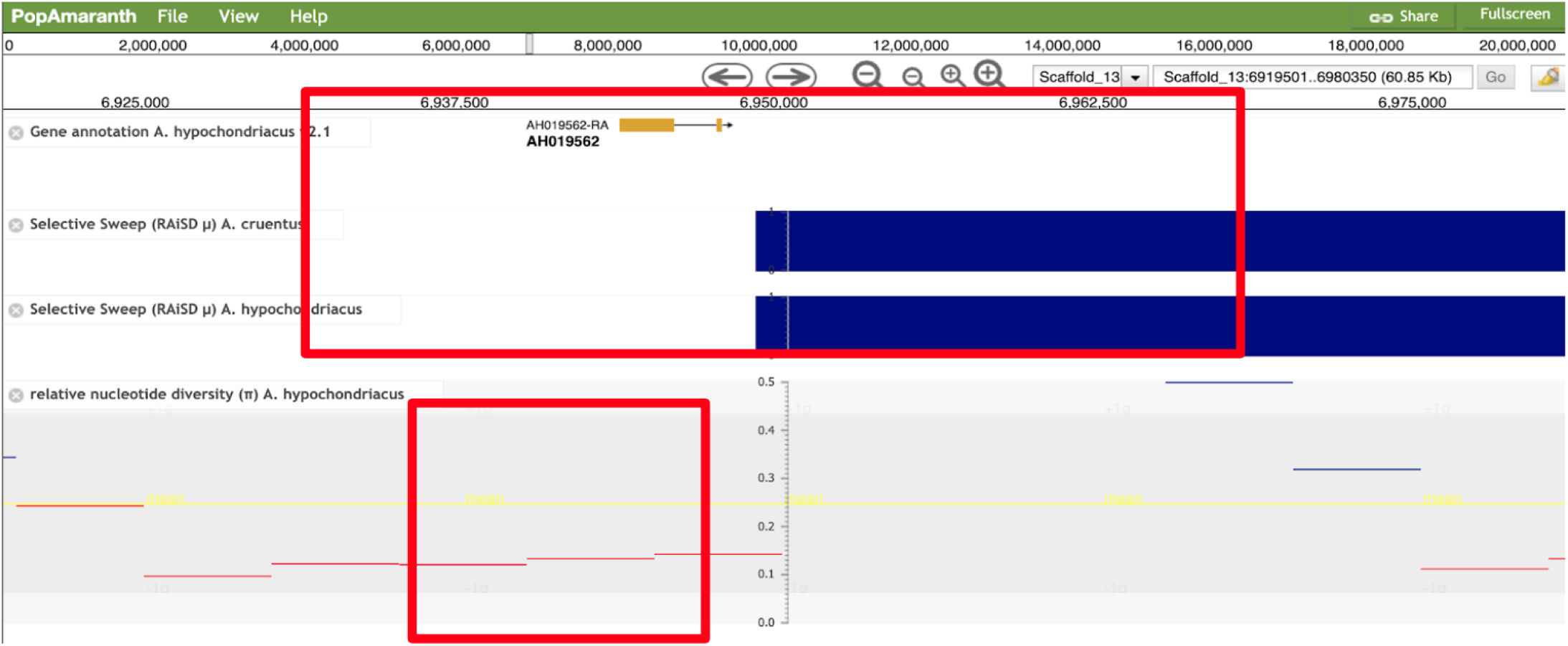
Screenshot of the gene AmTSAR1 (AH-019582) region. Signals of positive selection identified in *A. cruentus* and *A. hypochondriacus*. Relative nucleotide diversity between A. hypochondriacus compared to its wild ancestor *A. hybridus* is lower than the genome wide relative diversity, which is an indicator of selection in this region.

**Table S1.**
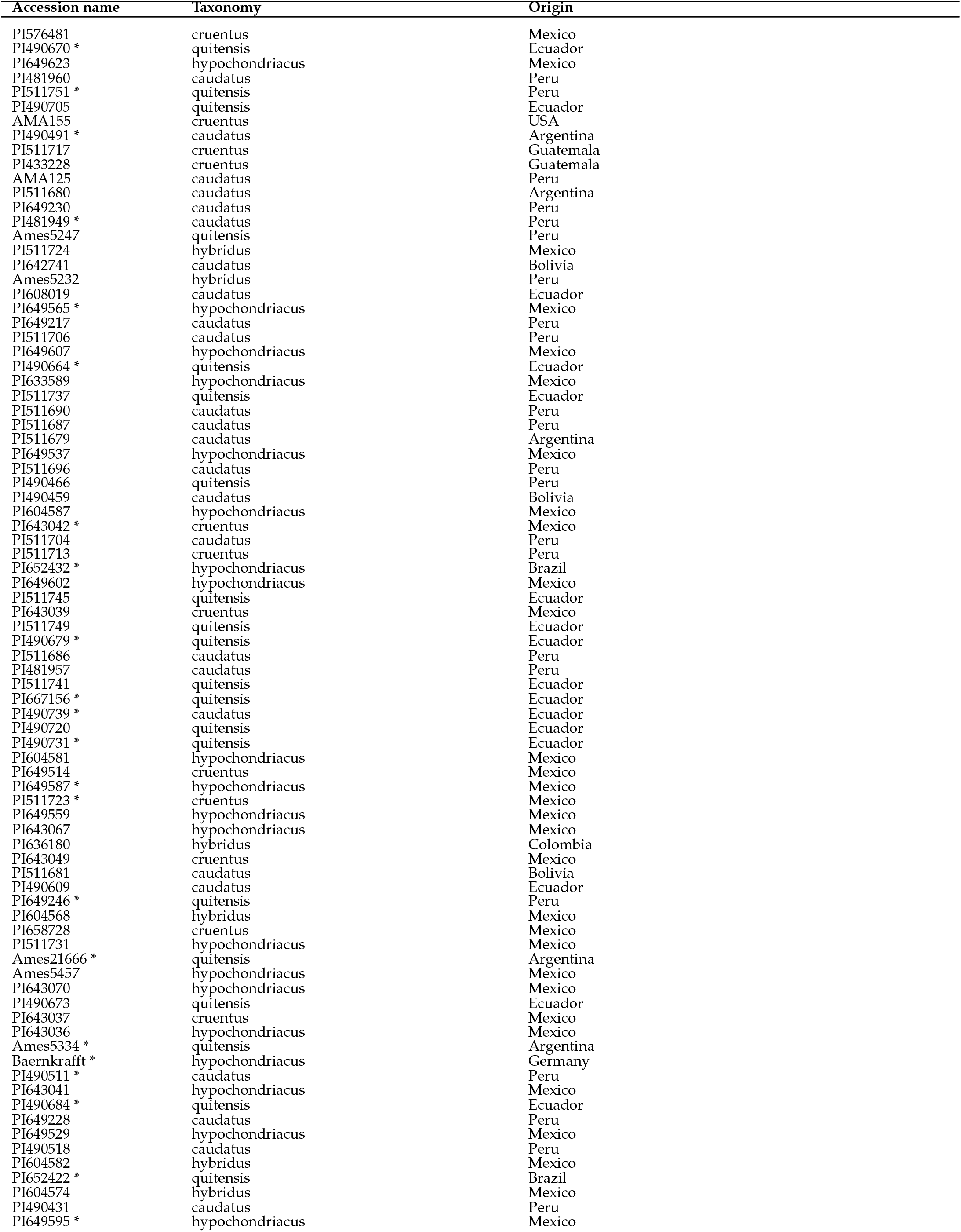
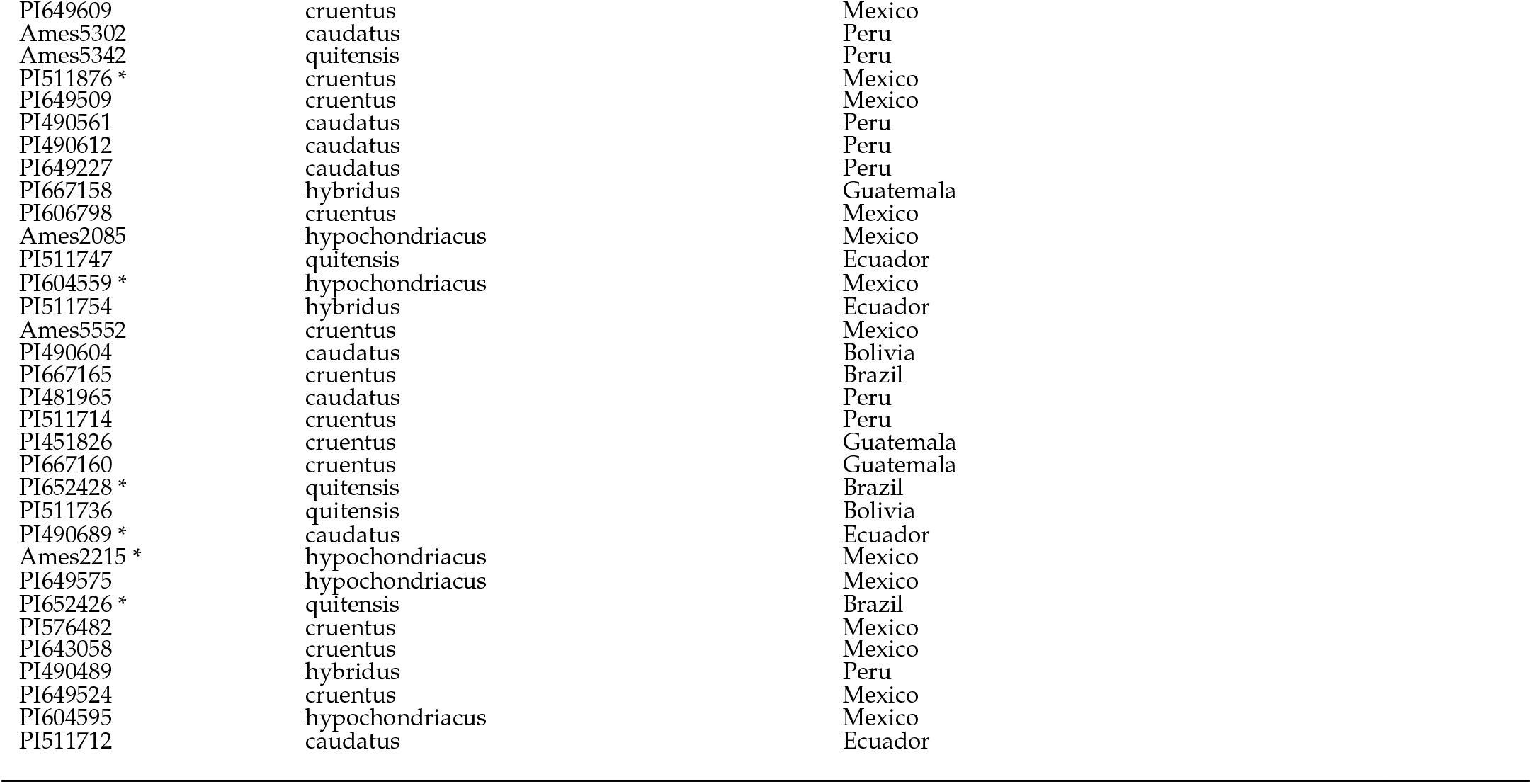
List of samples evaluated. Samples marked with * were filtered and not included in PopAmaranth

**Table S2.**
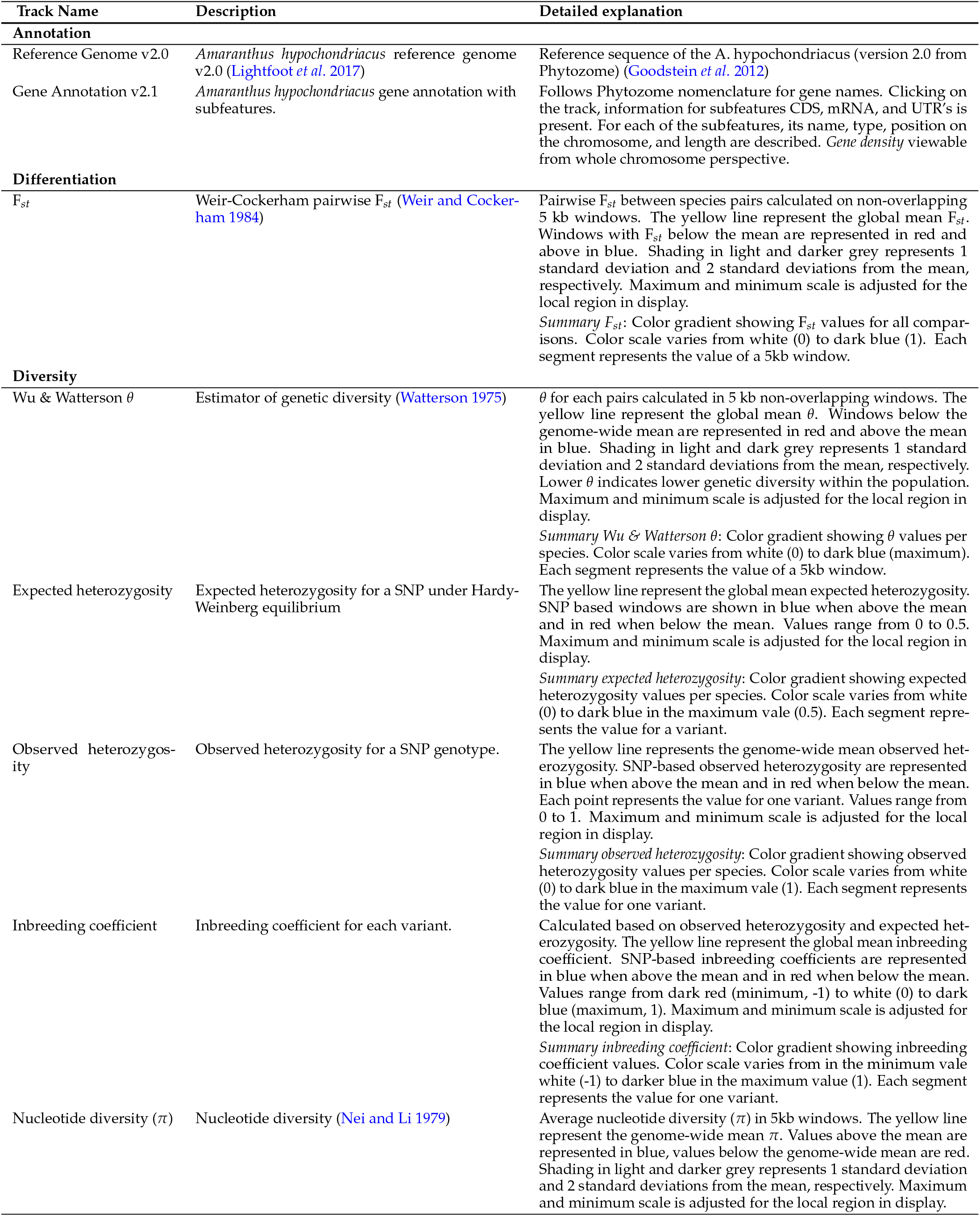
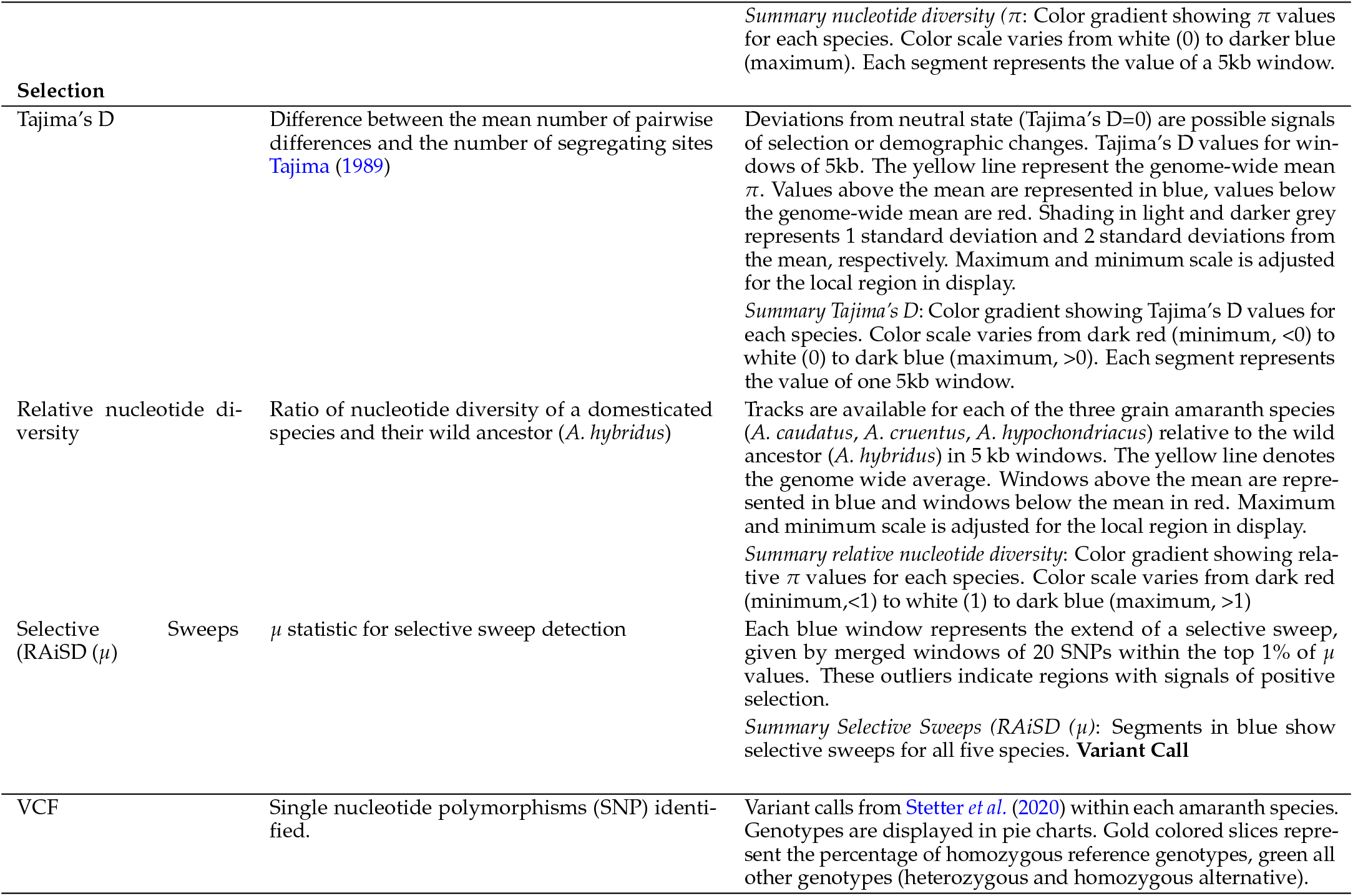
List of all tracks available on PopAmaranth at the time of publication. Detailed description of the included categories (bold) and respective tracks and summary statistics.

